# Soil bacterial and fungal communities show within field heterogeneity that varies by land management and distance metric

**DOI:** 10.1101/2022.07.26.501510

**Authors:** Fiona M. Seaton, Rob I. Griffiths, Tim Goodall, I Lebron, Lisa R. Norton

## Abstract

Increasing interest in the use of microbial metrics to evaluate soil health raises the issue of how fine-scale heterogeneity can affect microbial community measurements. Here we analyse bacterial and fungal communities of over 100 soil samples across 17 pasture farms and evaluate beta diversity at different scales. We find large variation in microbial communities between different points in the same field, and if Aitchison distance is used we find that within-field variation is as high as between-farm variation. However, if Bray-Curtis or Jaccard distance are used this variation is partially explained by differences in soil pH and vegetation and is higher under mob grazing for fungi. Hence, field scale variation in microbial communities can impact the evaluation of soil health.

---

There is increasing pressure to manage agriculture sustainably for both food production and environmental health (Tilman et al., 2011; Amundson et al., 2015). Microbial community structure and activity are often suggested to be key determinants of soil health (Stone et al., 2016; Bünemann et al., 2018), yet our understanding of how to use microbial data to guide farm management is still lacking (Fierer et al., 2021). A major issue in using microbial data within soil health metrics is the variability of soil microbial communities over space and time. It is known that soil properties such as soil pH, organic matter and nutrient content can all show fine scale spatial variation across agricultural landscapes (Ball and Williams, 1968; Lark et al., 2004; Kariuki et al., 2009). Therefore, it might be expected that microbial communities show similar levels of variation. Here we compare the levels of within-field variation in microbial communities to variation between fields and farms in order to evaluate the impacts of fine-scale variation upon microbial communities and to assess effects of land management, plant and soil properties.

In summer 2019, 17 pasture farms from across Great Britain within the Pasture Fed Livestock Association were surveyed for soil and vegetation properties. On each farm at least two fields undergoing differing land management practices were surveyed and within each field three sites were sampled for vegetation plus soil microbiological analysis and pH in water. In total, 110 samples of soil and vegetation were taken over 38 fields. Soil physicochemical properties were measured on bulked auger samples taken across each field in a W pattern. Land management within each field was categorised into ley, mob grazing, rotational grazing and set stocking based on farmer interviews. Bacterial and fungal communities were analysed through DNA sequencing of the 16S and ITS2 regions respectively. For a full description of the methods see Seaton et al. (2022). DNA sequences are publicly available in the European Nucleotide Archive under primary accession code PRJEB46195, sample accession codes ERS7103117 to ERS7103228. All statistical analysis was performed in R using the vegan and nlme packages (Oksanen et al., 2020; R Core Team, 2020; Pinheiro et al., 2021).

Comparison of the differences between microbial communities within fields indicated that while on average there was a gradient of increasing dissimilarity from within-field to within-farm to between-farm comparisons, within many fields the bacterial and fungal communities at different locations were as different to each other as they were to communities from different farms (Figure 1). Across both bacteria and fungi at least 80% of comparisons within field communities were within the 5-95% range of between field/within farm distances, and at least 40% of those within field comparisons were within the 5-95% range of between farm comparisons. Using a community dissimilarity metric that is suggested to be particularly effective at finding biological differences, i.e. Aitchison distance (Martino et al., 2019), microbial communities within the same field were on average as dissimilar as communities from different farms (Figure 1, bottom). In contrast, use of the Bray-Curtis or Jaccard metrics indicated that communities were more dissimilar between farms and fields than within fields. It is worth noting that the Aitchison distance involves weighting change in low abundance species equally to change in high abundance taxa while Bray-Curtis distance more strongly weights change in high abundance taxa and Jaccard distance ignores abundance. In this context abundance is based upon standardised read count, which is not necessarily related to biomass (Knight et al., 2018). The importance of rare taxa in determining ecosystem function is a much debated topic (Jousset et al., 2017). Our results show how conclusions can vary drastically depending on how strongly the rarer members of the community are weighted in the analysis.

**Figure 1:**
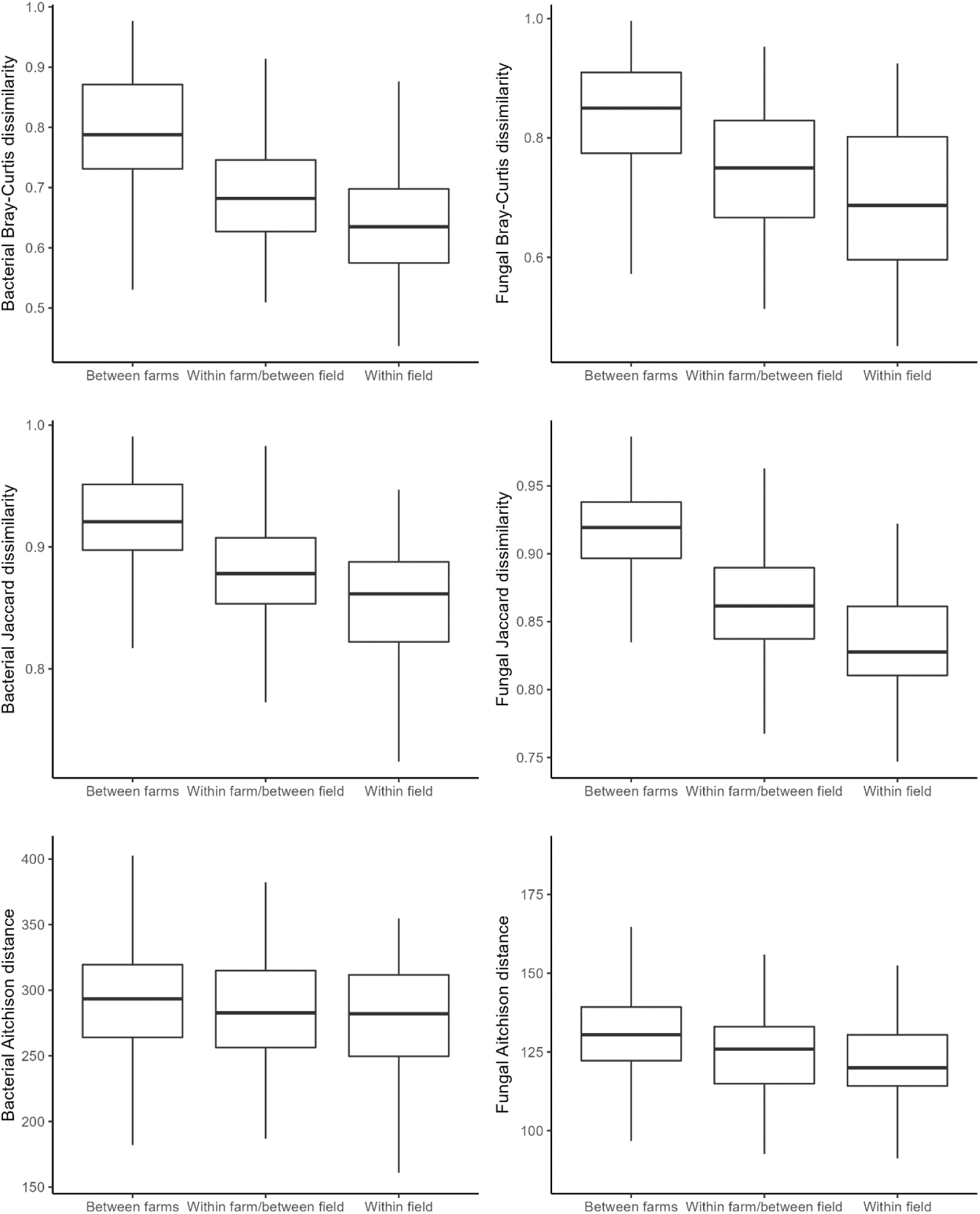
The community dissimilarity between farms, between different fields of the same farm, and between different points in the same field for bacteria (left) and fungi (right) as measured by Bray-Curtis distance (top), Jaccard distance (centre) and Aitchison distance (bottom).

Land management and plant community structure explained some of the differences between fungal communities at sampling locations on the same field and farm, but less so for bacterial communities. Mob and rotational grazing, which result in variable stocking levels across a field over time, resulted in somewhat higher variation in fungi than either set stocking or leys, but not in bacteria (Figure 2). Both the bacterial and fungal communities were strongly affected by the physicochemical soil variables, with soil pH and calcite explaining the first axis of variation and aggregate stability being roughly orthogonal (Table 1, Figures S1-2). The pH of the microbial sample showed a stronger relationship with community composition than the field average, showing the importance of including fine-scale edaphic information when evaluating responses of soil microbial communities to larger-scale drivers. Fungal communities showed greater associations with the plant community composition at each site, with a clear relationship with grass species richness (Table 1). The greater impacts of farm management and plant community composition upon fungi relative to bacteria is in agreement with previous studies of the soil microbial communities in British pasture (Seaton et al., 2022) and the known greater sensitivity of fungi to farm management techniques (Maharning et al., 2009).

**Table 1:** The variation in bacterial and fungal communities explained by the different plant and soil properties. Values are from fitting environmental data to NMDS ordinations of bacterial and fungal communities based on Bray-Curtis, binary Jaccard or Aitchison distance, information on the ordinations given in Figures S1-2. Soil pH is given for both the bulked samples across the field which were measured in CaCl_2_ and the pH in water for the microbial sample after freezing. Note that once the farm and field structure was accounted for there were no significant differences between the physicochemical or plant variables between land use types (all FDR-corrected p-values > 0.1). P values are not included as the soil variables are represented by one bulked sample measurement per field leading to inflated type I error. The plant ordination (PCoA) results are shown in Figure S3.

| Variable | Depth (cm) | Bacteria R<sup>2</sup> |  |  | Fungi R<sup>2</sup> |  |  |
| --- | --- | --- | --- | --- | --- | --- | --- |
|  |  | Bray-Curtis | Jaccard | Aitch. | Bray-Curtis | Jaccard | Aitch. |
| pH (micr. sample) | - | 0.856 | 0.861 | 0.443 | 0.728 | 0.721 | 0.545 |
| pH (CaCl<sub>2</sub>) | 0-5 | 0.602 | 0.672 | 0.498 | 0.508 | 0.526 | 0.335 |
|  | 5-15 | 0.716 | 0.764 | 0.548 | 0.571 | 0.582 | 0.424 |
| Electrical conductivity | 0-5 | 0.299 | 0.515 | 0.453 | 0.293 | 0.301 | 0.248 |
|  | 5-15 | 0.572 | 0.688 | 0.527 | 0.500 | 0.525 | 0.456 |
| Loss On Ignition (%) | 0-5 | 0.181 | 0.312 | 0.201 | 0.263 | 0.309 | 0.298 |
|  | 5-15 | 0.272 | 0.336 | 0.248 | 0.252 | 0.301 | 0.262 |
| Calcite | 0-5 | 0.323 | 0.370 | 0.375 | 0.248 | 0.275 | 0.438 |
|  | 5-15 | 0.310 | 0.365 | 0.344 | 0.248 | 0.278 | 0.439 |
| Clay | 0-5 | 0.278 | 0.305 | 0.143 | 0.102 | 0.143 | 0.130 |
|  | 5-15 | 0.365 | 0.396 | 0.200 | 0.181 | 0.233 | 0.176 |
| Silt | 0-5 | 0.057 | 0.097 | 0.068 | 0.110 | 0.115 | 0.066 |
|  | 5-15 | 0.031 | 0.061 | 0.055 | 0.057 | 0.064 | 0.063 |
| Sand | 0-5 | 0.067 | 0.080 | 0.040 | 0.061 | 0.060 | 0.056 |
|  | 5-15 | 0.070 | 0.086 | 0.055 | 0.185 | 0.041 | 0.077 |
| Aggregate stability | 0-5 | 0.140 | 0.175 | 0.286 | 0.132 | 0.135 | 0.086 |
|  | 5-15 | 0.189 | 0.215 | 0.088 | 0.173 | 0.199 | 0.131 |
| Forb richness | - | 0.104 | 0.158 | 0.100 | 0.034 | 0.020 | 0.239 |
| Grass richness | - | 0.101 | 0.147 | 0.137 | 0.403 | 0.370 | 0.250 |
| Plant PCoA 1 | - | 0.037 | 0.064 | 0.103 | 0.180 | 0.160 | 0.184 |
| Plant PCoA 2 | - | 0.172 | 0.182 | 0.213 | 0.098 | 0.101 | 0.094 |
| Plant PCoA 3 | - | 0.067 | 0.064 | 0.029 | 0.195 | 0.194 | 0.092 |
| Plant PCoA 4 | - | 0.128 | 0.141 | 0.043 | 0.135 | 0.146 | 0.257 |

**Figure 2:**
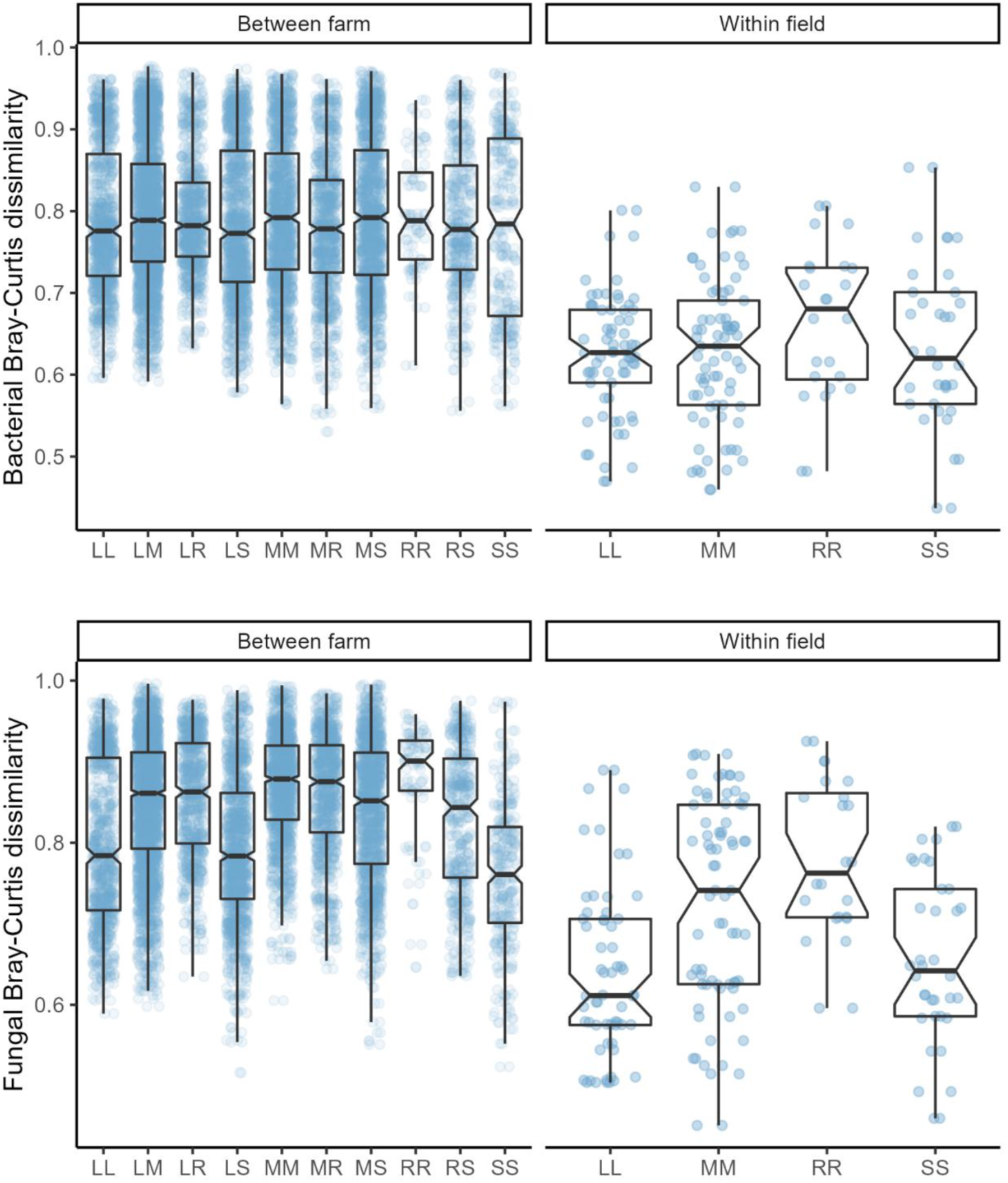
The Bray Curtis dissimilarity between farms and between different points in the same field, by land management technique for bacteria (top) and fungi (bottom). L indicates ley, M indicates mob grazing, R indicates rotational grazing and S indicates set stocking. Two of the same letter indicate a comparison between areas of the same land management, and two different letters indicate a comparison between the two land management types, e.g. LM indicates a comparison between ley and mob grazing.

Our results show that field-scale variability in soil microbial communities is widespread even in relatively homogenous land uses. The spatial scale at which we found high levels of heterogeneity is finer than that identified by previous studies of microbial community variation (Ranjard et al., 2013; Constancias et al., 2015). The standard approach to deal with high levels of within field variability when evaluating soil condition is to take several samples and then bulk them together (Kariuki et al., 2009). However, not only does the very small amount of soil used within standard DNA metabarcoding methods mean that a bulking approach will be unlikely to yield a true average of the field microbial community but also the compositional nature of DNA analyses means that the microbial community will not be exhaustively surveyed (Gloor et al., 2017). Therefore, the microbial community obtained within any one analysis is more likely to be either from an extreme environment relative to averaged soil properties and/or composed of multiple sections of communities that are in fact not co-located in reality. Our results also demonstrate that the patterns of variation are dependent on current land management and how we choose to evaluate them, i.e. which taxa are of interest, with mob grazing and rarer taxa showing greater variation at finer spatial scales. In summary, our results illustrate important considerations in the development and use of soil microbiome based characterisations of soil condition and show the importance of accounting for fine-scale variation within farm-level soil health assessment through survey design and inclusion of fine-scale soils information.

## Acknowledgements

We thank the PFLA and all the farmers involved in the survey described for working with us on this project. This work was supported by the Global Food Security Programme funded by UKRI (UK Research and Innovation), grant number (06211).

## Supplementary Figures

**Figure S1:**
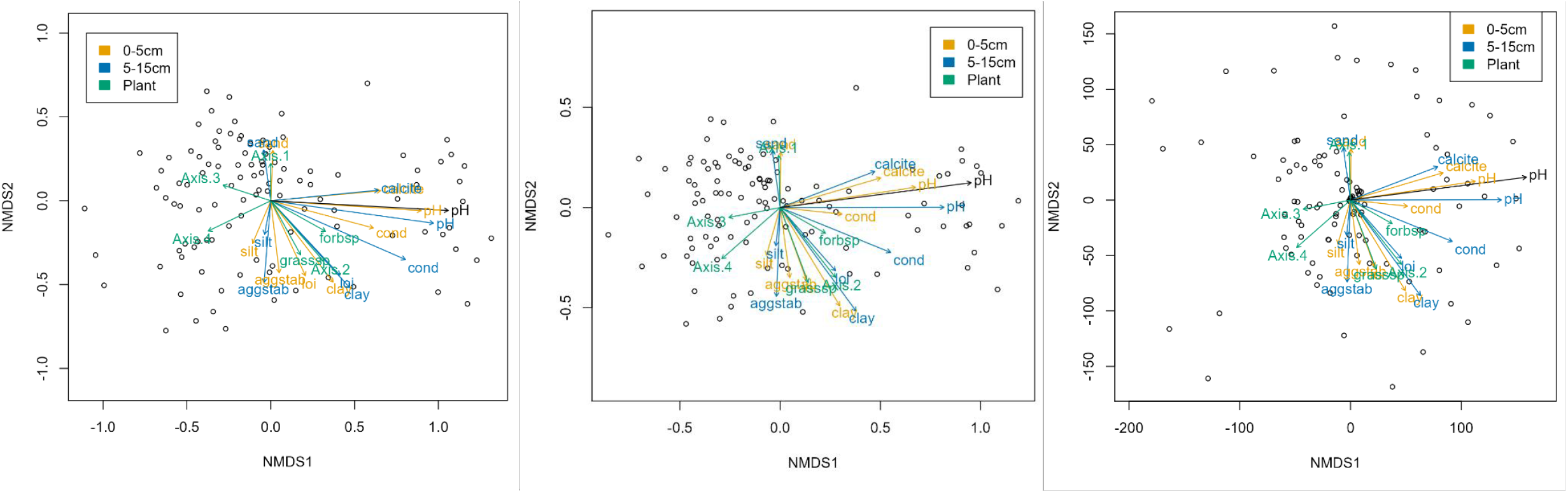
The first two axes of Bacterial NMDS ordination with Bray-Curtis distance (left), Jaccard distance (middle) and Aitchison distance (right). Stress values were 0.144 over two dimensions, 0.106 over three dimensions and 0.177 over four dimensions respectively. Ordinations using either Jaccard or Aitchison distance could not converge with only two dimensions included. Sites are represented by black circles and the fit of environmental vectors upon the NMDS scores are shown in coloured arrows. Variables included are pH (pH), electrical conductivity (cond), calcite (calcite), clay content (clay), silt content (silt), sand content (sand), soil organic matter/loss on ignition (loi), grass species richness (grasssp), forb species richness (forbsp), and the four plant PCoA axes (Axis.1, Axis.2, Axis.3 and Axis.4). Surface (0-5 cm) soil property measurements are shown in yellow and slightly deeper (5-15 cm) measurements in blue. The pH measurement shown in black represents the pH of microbial samples after freezing.

**Figure S2:**
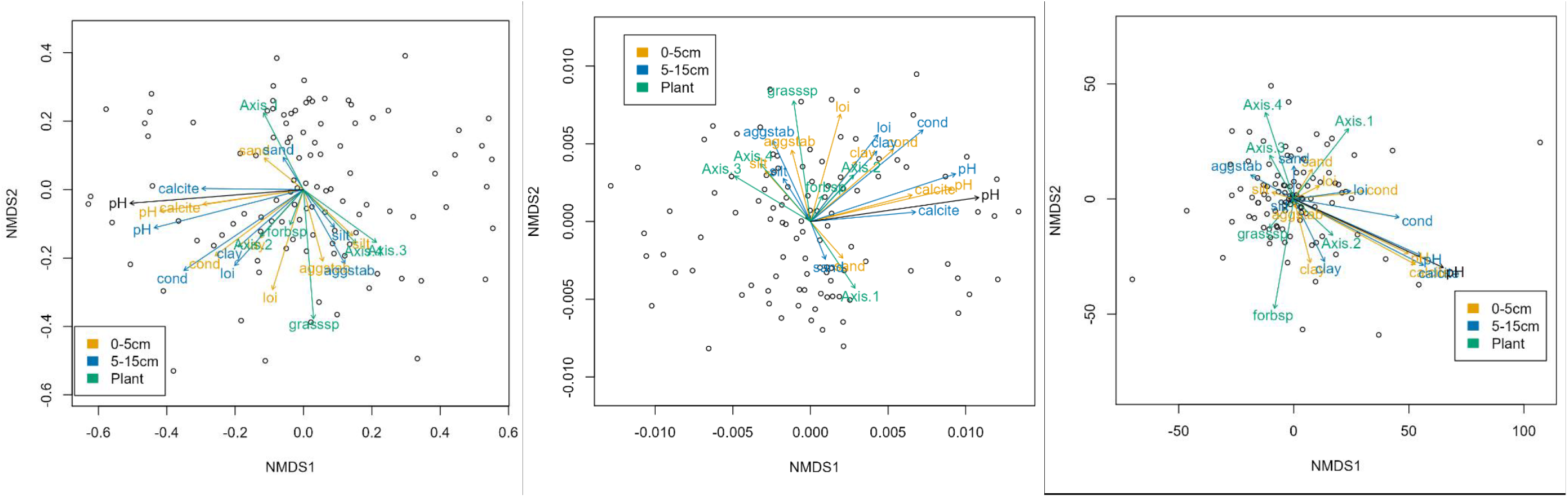
The first two axes of the fungal NMDS ordination with Bray-Curtis distance (left), Jaccard distance (middle) and Aitchison distance (right). Stress values were 0.204 over two dimensions, 0.209 over two dimensions and 0.207 over three dimensions respectively. NMDS using Aitchison distance could not converge with only two dimensions included. Sites are represented by black circles and the fit of environmental vectors upon the NMDS scores are shown in coloured arrows. Variables included are pH in CaCl2 (pH), electrical conductivity (cond), calcite (calcite), clay content (clay), silt content (silt), sand content (sand), soil organic matter/loss on ignition (loi), grass species richness (grasssp), forb species richness (forbsp), and the four plant PCoA axes (Axis.1, Axis.2, Axis.3 and Axis.4). Surface (0-5 cm) soil property measurements are shown in yellow and slightly deeper (5-15 cm) measurements in blue. The pH measurement shown in black represents the pH of microbial samples after freezing.

**Figure S3:**
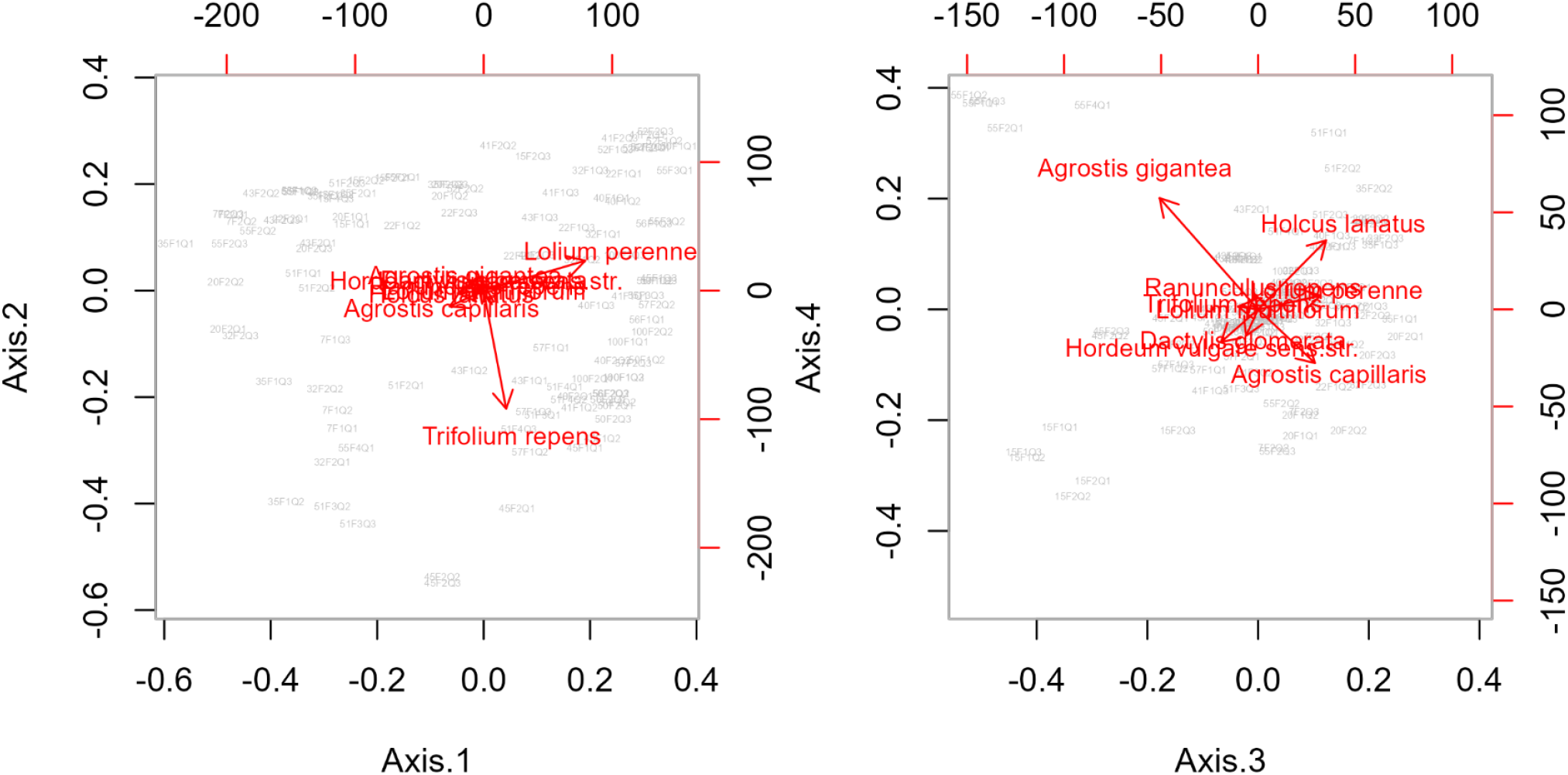
PCoA ordination results for plant community composition. Sites are represented by grey text, and plant species scores are shown as red arrows. Relative eigenvalues for Axis 1 through 4 were 0.289, 0.162, 0.105 and 0.074 respectively, or after Lingoes correction 0.092, 0.054, 0.037 and 0.028.

